# Low seroprevalence of neutralizing antibodies to gorilla adenovirus 32 (GRAd32) in southern African populations supports evaluation of this vector platform for HIV vaccine development

**DOI:** 10.64898/2026.05.23.726423

**Authors:** Nonhlanhla N. Mkhize, Anna Patjane, Nomcebo Shusha, Ashleigh Welsh, Tandile Hermanus, Prudence Kgagudi, Thopisang Motlou, Linda-Gail Bekker, Glenda E. Gray, Nigel Garrett, Lee Fairlie, Alex Sigal, Wendy A. Burgers, Takudzwa Magwaku, Tariro Makadzange, Stefano Colloca, Antonella Folgori, Thandeka Moyo-Gwete, Michela Gentile, Stefania Capone, Penny L Moore

**Author notes:** Corresponding author: Dr Nonhlanhla N. Mkhize. Contributed equally (Nonhlanhla N. Mkhize, Anna Patjane). Contributed equally (Stefania Capone, Penny L Moore). Deceased (Thopisang Motlou).

## Abstract

Adenoviruses (Ads) are widely used as vaccine vectors. However, pre-existing immunity to commonly used serotypes, like Ad5, can reduce vaccine immunogenicity, with neutralizing antibody titers >200 previously shown to impact vaccine efficacy. Gorilla adenovirus (GRAd) vectors have been developed to evade pre-existing anti-vector responses, but their seroprevalence in southern Africa is poorly defined. Here, we assessed seroprevalence to GRAd32, Ad26 and Ad5 before (baseline) and after COVID-19 vaccination, in cohorts from South Africa and Zimbabwe.

Sera from South African participants enrolled in the Sisonke sub-study (n=100, prior to Ad26.COV2.S vaccination) and the follow-up “Booster after Sisonke” (BaSiS) study (n=226) were tested for neutralizing antibodies to Ad5, Ad26, and GRAd32. These samples included paired pre/post boost samples for 27 donors. We also tested sera from the Zimbabwean Mutala cohort (n=131, of which 44 were unvaccinated, and 87 vaccinated with inactivated vaccines). Participants living with HIV (PLWH) comprised 30-50% of each cohort.

In the pre-vaccination samples from the Sisonke cohort, geometric mean titers (GMT) for anti-GRAd32, Ad26, and Ad5 antibodies were 78, 142, and 459, with neutralization titers >200 observed in 14%, 32%, and 68% of participants, respectively. Similarly, in the unvaccinated participants in the Mutala cohort, GMTs for GRAd32, Ad26, and Ad5 were 117, 245, and 536, with neutralizing titers >200 in 22%, 42%, and 69% of participants. We observed no significant difference in Ad antibody titers between PLWH and those living without HIV. We next assessed the impact of COVID-19 vaccination on titers. Vaccination with inactivated COVID-19 vaccines (Sinopharm/Sinovac) did not significantly affect Ad5, Ad26 or GRAd32 titers in an unpaired analysis. In contrast, ∼9 months after Ad26.COV2.S vaccination, anti-Ad26 titers for longitudinally sampled participants (n=27) increased 10-fold from a GMT of 141 to 1,426. By comparison, GRAd32 responses were not significantly altered by Ad26.COV2.S vaccination, while anti-Ad5 responses showed a modest <2-fold increase.

Our data support previous findings that, whereas anti-Ad5 neutralizing antibody responses are commonly detected globally, GRAd32 responses are less frequent. Importantly, GRAd32 neutralizing responses remained unchanged after Ad26.COV2.S vaccination. HIV status had no significant effect on antibody titers. These results support the use of the GRAd32 vector in upcoming HIV vaccine trials, including in regions where Ad26-based COVID-19 vaccines were widely deployed.

## INTRODUCTION

Adenoviral (Ad) vectors are widely used in vaccines against major infectious diseases, including Ebola, HIV, and SARS-CoV-2 (Chang 2021; Buchbinder et al. 2008; Zhu, Guan, et al. 2020; Sayedahmed et al. 2020). Both human-derived adenoviruses (e.g., adenovirus type 5 (Ad5), adenovirus type 26 (Ad26)) and non-human primate vectors (e.g., chimpanzee adenovirus ChAdOx1 and gorilla adenovirus 32 (GRAd32)) have successfully induced cellular and humoral immunity (Zhu, Li, et al. 2020; Capone et al. 2023; Barouch et al. 2021, 2013; Ewer et al. 2021). In addition to the fact that vaccines in these vectors are immunogenic and have a good safety profile, Ad vectors are easy to manipulate, do not require specialized cold chain storage, and are generally cost effective (Coughlan et al. 2022). Indeed, their utility as an emergency vaccine platform was validated during the SARS-CoV-2 pandemic (Madhi et al. 2021; Bekker et al. 2022; Le Gars et al. 2024).

One potential drawback of viral-vectored vaccines is that pre-existing neutralizing antibodies may reduce their effectiveness. For Ad5, titers of 200-1,000 have been associated with reduced immunogenicity (Priddy et al. 2008). Seroepidemiological studies have consistently found a high prevalence of Ad5 neutralizing antibodies in Africa, Asia and South America (often >70%), with lower prevalence in Europe and North America (Barouch et al. 2011; Mennechet et al. 2019). Prior to the COVID-19 pandemic, Ad26-specific immunity was less common worldwide, in less than 10% of adults in developed regions, but in 31% and 66% of Brazilian and South African adults, respectively (Barouch et al. 2011; Fuchs et al. 2015). Despite this, pre-existing Ad26 neutralizing antibodies had a limited effect on immunogenicity for Ad26.COV2.S and Ad26.ZEBOV (McDonald et al. 2021; Le Gars et al. 2024). Similarly, while a failure of Ad26. COV2.S to boost antibody titers was reported in a South African trial, this could not be definitively attributed to anti-vector immunity (Riou et al. 2024).

Mass immunization with Ad26.COV2.S (Johnson & Johnson) and ChAdOx1 nCoV-19 (Oxford–AstraZeneca) COVID-19 vaccines was a central component of Africa’s vaccination response, especially during the initial rollout phases (Rees and Madhi 2021). In South Africa, the Sisonke trial alone vaccinated over 470,000 health-care workers in South Africa using Ad26.COV2.S (Bekker et al. 2022), though the Pfizer vaccine became increasingly used later in the pandemic. Zimbabwe, on the other hand, relied largely on Sinopharm and Sinovac inactivated vaccines, with adenoviral vaccines playing a relatively minor role in the country’s response to SARS-CoV-2. Whether these resulted in population-level increases in the prevalence of anti-adenovirus antibodies has not been described. Southern Africa also bears the brunt of the HIV pandemic (Kharsany and Karim 2016), which may impact baseline prevalence and titers of Ad5 and Ad26, though this has not been evaluated for GRAd32 (Luo et al. 2024; Sun et al. 2011). These findings reinforce the importance of selecting adenoviral vectors with minimal background immunity in African settings — both to optimize primary responses and to preserve boosting potential.

Neutralizing antibodies to simian adenoviruses derived from chimpanzees, bonobos, and gorillas are rare (0%–18%) in the human population (Capone et al. 2021; Quinn et al. 2013; Colloca et al.2012). For ChAdOx1 (chimpanzee adenovirus), studies found very low baseline prevalence of neutralizing antibodies in most populations, with little evidence of reduced immunogenicity due to pre-existing antibodies (Dicks et al. 2012; Hong et al. 2025; Xiang et al. 2006). Similarly for gorilla adenovirus 32 (GRAd32), Capone et al. reported low levels of nAbs in US and Italian serum cohorts (Capone et al. 2021).

On the basis of low seroprevalence in the US and Italy, and robust induction of SARS-CoV-2 cellular immunity (Capone et al. 2023; Capone et al. 2021; Lanini et al. 2022; Agrati et al. 2022), GRAd32 is being explored as a novel vaccine vector. This includes the HIV vaccine, Gorilla Adenovirus Vectored HIV Networked Epitopes Vaccine (GRAdHIVNE1) that is now being tested in a first-in-human phase 1 clinical trial, IAVI C114 (NCT06617091). GRAdHIVNE1 employs a GRAd vector to deliver a T cell immunogen composed of highly “networked” mutationally constrained regions of the HIV proteome — sites where structural connectivity and functional necessity limit viral escape. These conserved epitopes are frequently targeted in natural HIV control (Gaiha et al. 2019) and may elicit potent, broadly reactive CD8⁺ T cell responses selected for wide HLA coverage. This vaccine concept is now in early clinical testing in African populations, where it will be important to account for high baseline immunity to common adenoviral vectors.

We therefore evaluated the prevalence of neutralizing antibodies to GRAd32 in two southern African populations, placing these data in the broader context of adenoviral vector immunoepidemiology and the specific challenges of HIV-endemic settings. We show that potentially immune interfering titers (>200) to GRAd32 are detected in less than 20% of participants from South Africa and Zimbabwe, even after Ad26.COV2.S COVID-19 vaccination and in the context of HIV co-infection. These data support further evaluation of a GRAd-based HIV vaccine in African settings, where such a vaccine is most needed.

## METHODS

### Samples and ethical approvals

This seroprevalence study was approved by the Human Research Medical Ethics Committee at the University of the Witwatersrand (ethics numbers M240570 and M240810). All participants provided written informed consent. Samples were tested from four groups, two from South Africa and two from Zimbabwe, which had varying exposure to COVID-19 vaccines (**Table 1**), and compared to commercial samples from the United States of America.

**Table 1:**
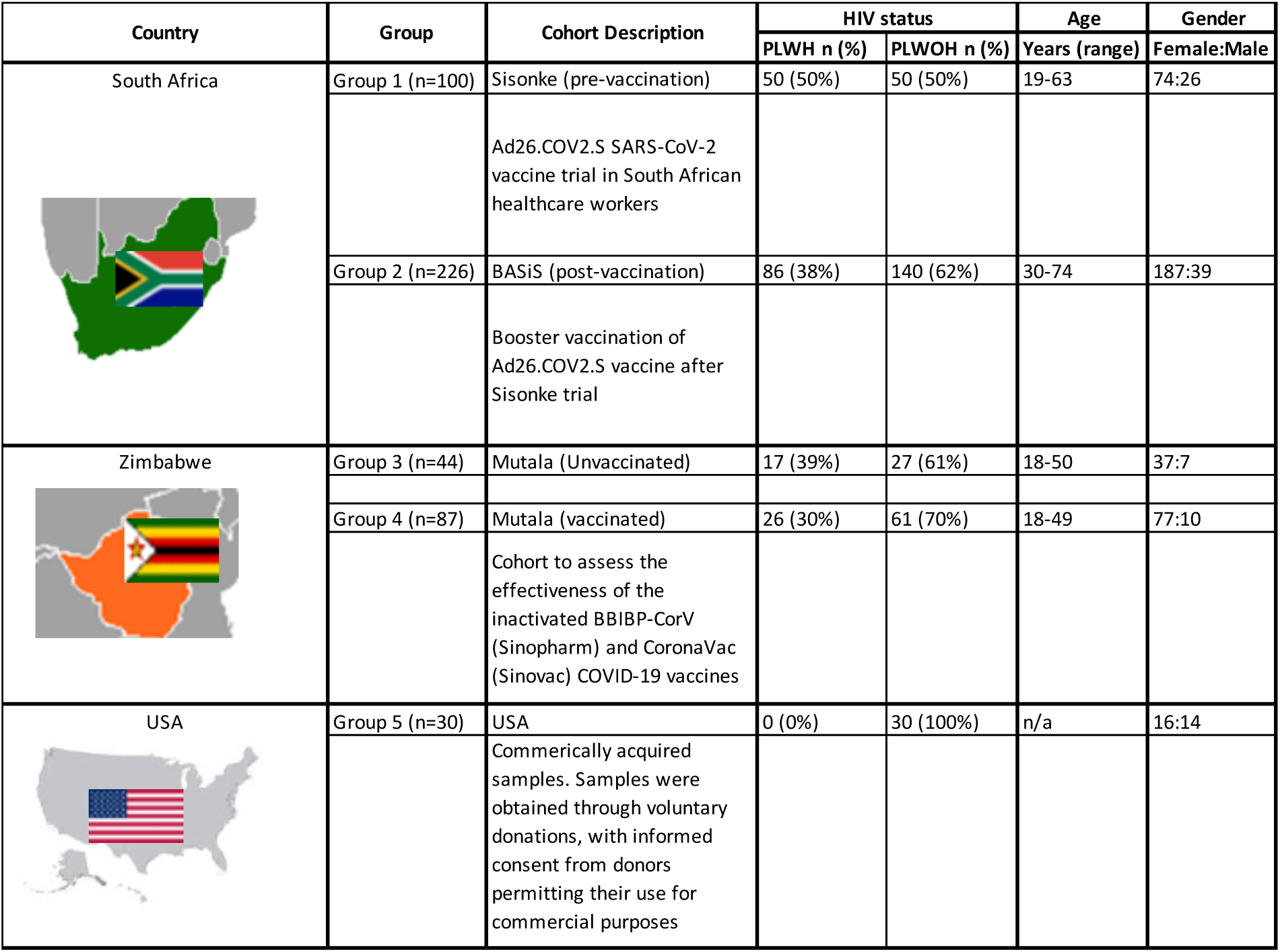
Demographic data of the South African, Zimbabwean and USA Cohorts.

Group 1 consisted of baseline (i.e. pre-vaccination) samples from 100 South African participants enrolled in an immunogenicity substudy of the Sisonke clinical trial of the Ad26.COV2.S vaccine in healthcare workers (HCWs). The Sisonke trial was a single-arm, open-label, phase 3B implementation study of the Ad26.COV2.S SARS-CoV-2 vaccine in South African healthcare workers aged 18 years and over. Participants were recruited between 17 February 2021 and 17 May 2021 and received a single dose of 5×10¹⁰ viral particles of the Ad26.COV2.S vaccine. Approximately 50% of this group were people living with HIV (PLWH), whose HIV infection was well controlled due to high levels of antiretroviral (ARV) usage (Bekker et al. 2022). This study was registered with the South African National Clinical Trial Registry (DOH-27-022021-6844), ClinicalTrials.gov (NCT04838795) and the Pan African Clinical Trials Registry (PACTR202102855526180) (Bekker et al. 2022).

Group 2 comprised samples from 226 South African participants who had taken part in the Sisonke trial described above and had subsequently been enrolled in the Booster After Sisonke Study (BaSiS) (Riou et al. 2024). The BaSiS trial was a randomised, open-label, phase 2 study designed to evaluate the safety and immunogenicity of a booster vaccination in participants who had received a single dose of the Ad26.COV2.S vaccine through the Sisonke phase 3B implementation study or via the South African National Department of Health’s rollout of the Pfizer-BioNTech BNT162b2 vaccine. These samples were analyzed at enrolment into BaSiS, approximately nine months following the Ad26.COV2.S vaccination received through Sisonke, between 12 August 2021 and 27 July 2022, to assess the impact of adenovirus-based SARS-CoV-2 vaccination on antibody titers. Of these, 38% were PLWH, who were generally accessing ART and virologically suppressed. This study was approved by the South African Health Products Regulatory Authority (SAHPRA, number: 20210423) and all site-specific Human Research Ethics Committees (Wits: 211001B, UKZN: BREC/00003487/2021, UCT: 680/202) (Riou et al. 2024). For a small subset (n = 27), paired samples were available at enrolment in both Sisonke and BaSiS and were assessed longitudinally to evaluate the impact of vaccination.

Groups 3 and 4 comprised post-pandemic samples obtained from 131 adults recruited by the Mutala site in Harare, Zimbabwe, between 2021 and 2022. This study was approved by the Medical Research Council of Zimbabwe (MRCZ) (number: MRCZ/A/2914). Of these, 44 participants had not received a vaccine for SARS-CoV-2 (Group 3), whereas 87 participants had been vaccinated with inactivated Sinovac and Sinopharm vaccines (Group 4). None of the Group 4 participants reported being vaccinated with adenoviral vaccines. There were 39% PLWH in group 3 and 30% PLWH in group 4.

Group 5 comprised pre-pandemic human serum samples from unvaccinated participants (n = 30; 14 male and 16 female) recruited in the United States of America and procured from TCS Biosciences (Botolph Claydon, Buckingham, UK) as controls. The samples were obtained through voluntary donations, with informed consent from donors permitting their use for commercial purposes. Donor confidentiality was maintained through strict data protection measures, including secure data handling and complete delinking of identifying information. The supplier is registered with the United States Food and Drug Administration (FDA) as a Blood Establishment (Capone et al. 2021). These samples were sourced prior to the start of the pandemic, and the participants had not been vaccinated against COVID-19.

### Neutralization assay

NAb titers in human sera were assayed as previously described (Capone et al. 2021; Aste-Amézaga et al. 2004). Briefly, 8 × 10^4^ HEK293 cells per well were seeded in 96-well plates the day before the assay. Each adenoviral vector (GRAd32, Ad5 or Ad26) encoding for secreted alkaline phosphatase (SEAP) was preincubated for 1 h at 37°C alone or with serial dilutions of control or test serum samples and then added to the 80%–90% confluent HEK293 cells. Genetically engineered Adenovirus lysate stocks of GRAd32, Ad5 or Ad26 expressing the gene for secreted alkaline phosphatase (SEAP) were received from ReiThera Srl (Rome, Italy), with infectious titers determined in the lab using the tissue infectious dose 50% (TCID_50_) assays and used at a multiplicity of infection (MOI) of 6 for GRAd32, 30 for Ad26 and 0,7 for Ad5. After incubation for 1 h at 37°C, supernatant was then removed and replaced with 10% FBS in DMEM. SEAP activity was measured 24 h later with the chemiluminescent substrate from the Phospha-Light kit (Applied Biosystems, Foster City, CA, USA). Neutralization titers were defined as the dilution at which a 50% reduction of SEAP activity from serum sample was observed relative to SEAP activity from virus alone. The neutralization titer, expressed as an ID_50_ value, was calculated using the nonlinear regression—[inhibitor] vs. normalized response variable slope method—by using the GraphPad Prism 8.0.1 software.

### Statistical analysis

The Wilcoxon matched-pairs signed rank test was used to determine statistical significance between baseline and nine months post-vaccination time points. Spearman’s non-parametric test was used to calculate the correlation between anti-Ad responses across all serotypes. Geometric mean values and fold changes are shown under the relevant graphs. Analyses were conducted using GraphPad Prism v10.0.2. Significance is shown as: ***<0.001, **p<0.01, *p<0.05.

## RESULTS

### Neutralizing antibodies to GRAd32 are rare in unvaccinated South African and Zimbabwean adults

Neutralizing antibodies to GRAd32 were demonstrated to be rare in a small study of participants from Italy, with only 11% of samples having titers greater than 200 (Capone et al. 2021). Here, we extended these studies to assess GRAd32 seroprevalence in COVID-19 unvaccinated South African (n=100, Group 1) and Zimbabwean (n=44, Group 3) cohorts.

We observed low levels of neutralizing antibodies to GRAd32 in both unvaccinated African cohorts, with geometric mean titers (GMT) ranging from 78 (South Africa) to 117 (Zimbabwe) (**Fig 1A**). There were no detectable responses to GRAd32 (determined as titers <50) in 58% of South African participants, and in 39% of Zimbabwean samples (**Fig 1B**). Furthermore, only 14% of South Africans and 22% of Zimbabweans had GRAd32 nAb responses >200, which is comparable to those in the USA (Group 5) (GMT of 67, with 13% of participants having titers >200, **S1 Fig Fig 1B**), and to titers previously reported in Italy (Capone et al. 2021; Lanini et al. 2022). A similar analysis of Ad26 and Ad5 in unvaccinated South African and Zimbabwean adults revealed higher levels of seropositivity. For Ad26, GMT ranged from 142 in unvaccinated South African adults to 245 in unvaccinated Zimbabwean adults **(Fig 1C)**, with 32% and 42% of participants in South Africa and Zimbabwe, respectively, having titers >200 **(Fig 1D)**. For Ad5, as expected, even higher titers were observed (GMT of 459 and 536, in South African and Zimbabwean adults, respectively) **(Fig 1E)**. Most participants (>68%), regardless of geographic location, had titers exceeding 1:200 **(Fig 1F)**. Overall, while Ad26 and Ad5 levels were similar to those reported in previous studies, the rare detection of GRAd32 in unvaccinated adults in South Africa and Zimbabwe supports the use of GRAdHIVNE1 in these populations.

**Fig 1:**
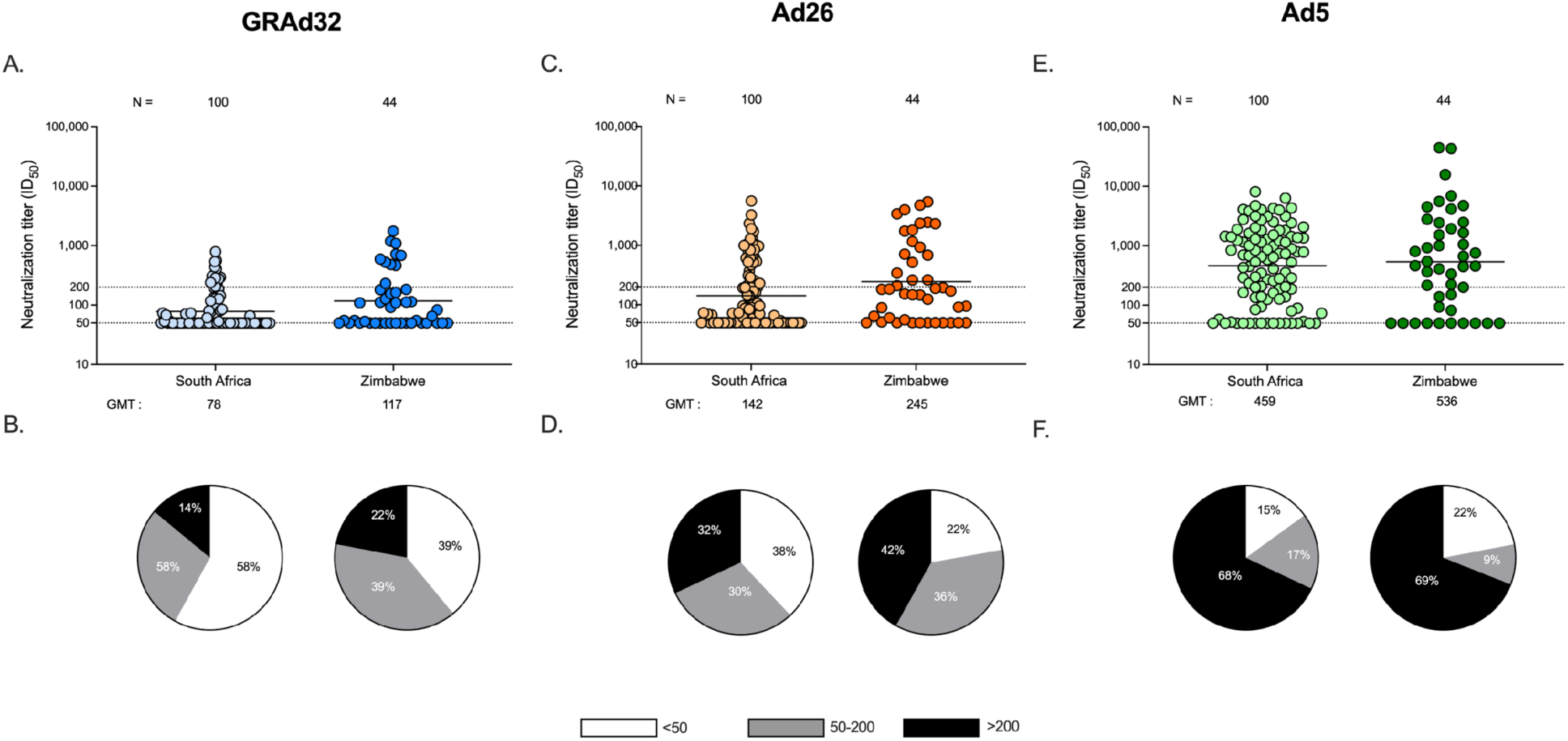
Neutralizing antibodies to GRAd32 are rare in unvaccinated South African and Zimbabwean adults. Neutralizing antibody responses to GRAd32 (A and B), Ad26 (C and D) and Ad5 (E and F) in COVID-19 unvaccinated adults from South Africa and Zimbabwe. In panels A, C and E, each dot represents a single participant. Geometric mean titers (GMT) are shown by horizontal lines, and values are shown below the graph. Dotted lines indicate titers of 50 (the lower limit of detection) and 200 (a titer above which reduced immunogenicity has been reported). In panels B, D and E, distribution of neutralization antibody titers to GRAd32 (B), Ad26 (D) and Ad5 (F) from each of the cohorts with proportions of titers <50 (white), titers between 50 and 200 (gray) and titers >200 (black) shown in the form of pie charts.

### Ad26.COV2.S vaccination, but not inactivated SARS-CoV-2 vaccines, boosts titers to Ad26, but does not impact GRAd32 titers

We next compared the prevalence of neutralizing antibodies to GRAd32, Ad26 and Ad5, before and after COVID-19 vaccination. We first analyzed 27 sets of paired samples from participants enrolled into the Sisonke trial, and approximately 9 months later after a single Ad26.COV2.S vaccine, into the BaSiS trial. We observed no significant increase in GRAd32 titers (GMT of 55 pre-vaccination vs a GMT of 61, 9 months post-vaccination) (**Fig 2A**). We observed a modest but statistically significant 1,7 fold increase in Ad5 titers from 405 to 691 (p=0.0002) (**Fig 2A**), which may suggest some degree of cross-reactivity of boosted Ad26 titers for Ad5. In contrast, Ad26 titers were significantly boosted by 10-fold from a GMT of 141 to 1,426 (p<0.0001) (**Fig 2A**).

**Fig 2:**
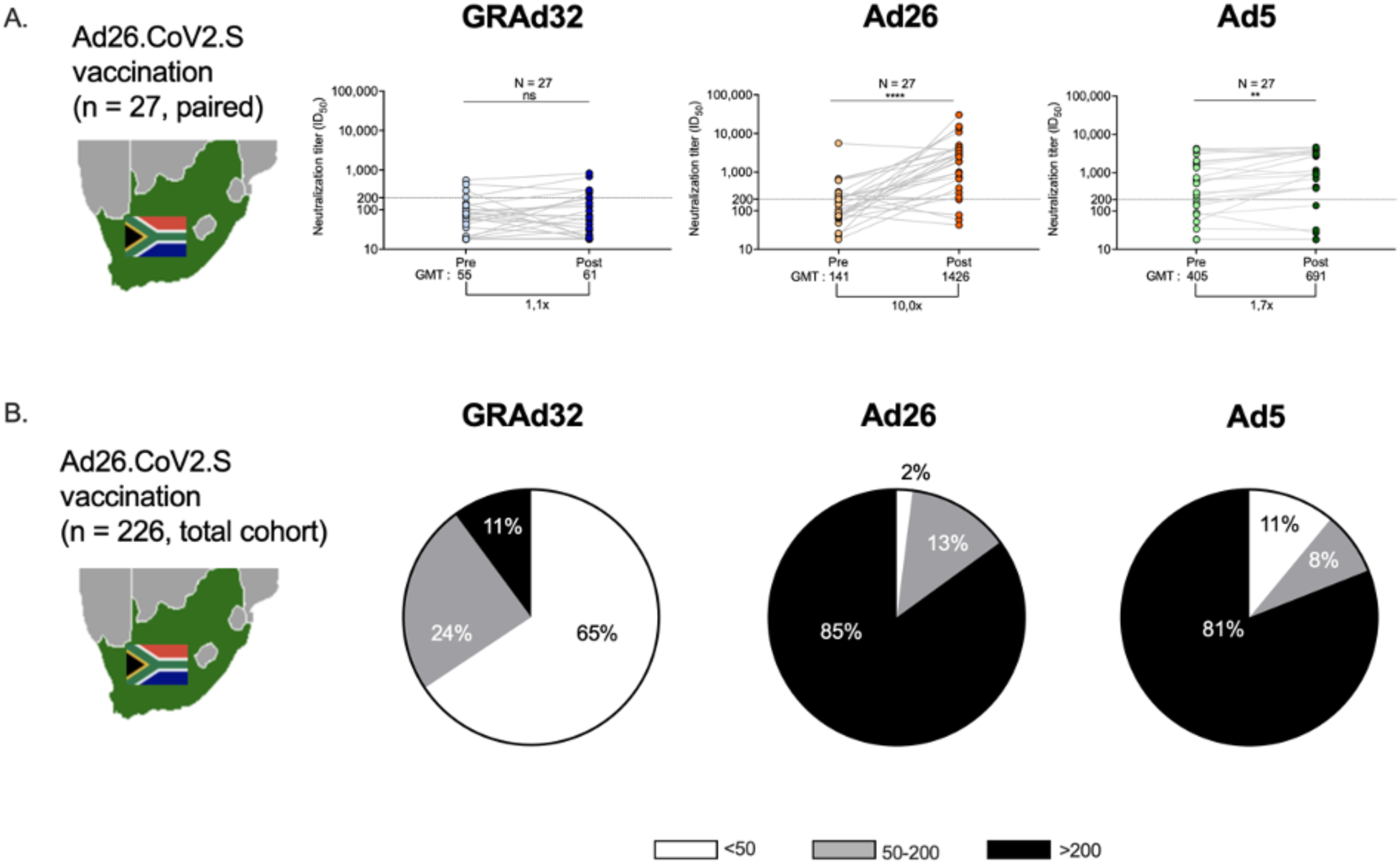
Ad26.COV2.S vaccination results in increased titers to Ad26 and Ad5, but does not impact GRAd32 titers. **A**) Paired samples from individuals who were enrolled into Sisonke and later into BaSiS, 9 months after Ad26.COV2.S vaccination, were tested for anti-Ad responses: GRAd32 (blue); Ad26 (orange); Ad5 (green). Each dot represents a single participant. Geometric mean titers (GMT) are shown by horizontal lines, and values are shown below the graph, together with fold changes. Dotted line indicate titers of 200 (titer associated with reduced immunogenicity for vaccines). **B**) Pie-charts showing the distribution of Ad titers in the unpaired cohort of individuals (n=226) enrolled in the BaSiS cohort (after vaccination).

To assess post-vaccination responses in a larger, albeit unpaired, cohort, we measured nAbs to all three adenoviruses in all 226 participants as they were enrolled into the BaSiS cohort (**Fig 2B)**. For GRAd32, titers remained stable and low (GMT of 74, compared to 78 before vaccination) (**S2 Fig A and B**), with only 11% having titers >200 (**Fig 2B**). In contrast, as with the small, paired cohort, titers to Ad26 were significantly higher post-vaccination with the GMT going from 142 to 1,509 (**S2 Fig A and B**), and 85% of participants having titers >200 (**Fig 2B**). Also as seen in the paired analysis, titers to Ad5 were increased, going from a GMT of 459 to 960 (**S2 Fig A and B**), with a concomitant increase in the proportion of participants having titers >200, to 81% (**Fig 2B**). This suggests that having higher Ad26 titers does not in itself result in increased cross-reactivity for GRAd32, and indeed we observed no correlation between Ad26 and GRAd32 titers, between Ad5 and GRAd32 titers, or between Ad5 and Ad26 titers (data not shown).

Lastly, we confirmed that boosting of Ad26 titers in the South African cohort was specifically due to exposure to Ad26.COV2.S, by assessing Ad26 nAb titers in Zimbabwean participants vaccinated with inactivated vaccines (Sinovac, n=33 and Sinopharm, n=54, compared to unvaccinated individuals (**S3 Fig A-C**). In these participants, titers to GRAd32, Ad26 and Ad5 were similar and there was no significant difference when compared to the unvaccinated group (**S3 Fig).** Thus, we conclude that vaccination with Ad26.COV2.S, but not inactivated vaccines, significantly increases Ad26 titers, slightly boosts Ad5 titers, but does not impact GRAd32 titers.

### Titers to GRAd32 and other vaccine-relevant adenoviruses are similar in PLWH and PLWOH

Lastly, we assessed whether anti-adenovirus nAbs differed in PLWH compared to people living without HIV (PLWOH). In the unvaccinated Sisonke cohort, where 50% (n=50) of participants were PLWH, we compared nAb titers to GRAd32, Ad5 and Ad26 between the two groups (PLWH and PLWOH) **(Fig 3A and B**). We observed no significant differences either in terms of GMT or proportion of donors with titers >200. Similarly, for the unvaccinated Zimbabwean cohort, titers in PLWH did not differ from those not living with HIV (**Fig 3C and D**). There were also no differences observed between vaccinated PLWH and PLWOH in South Africa and Zimbabwe (data not shown). Overall, HIV status did not affect adenovirus nAb responses in individuals from both South Africa and Zimbabwe.

**Fig 3.**
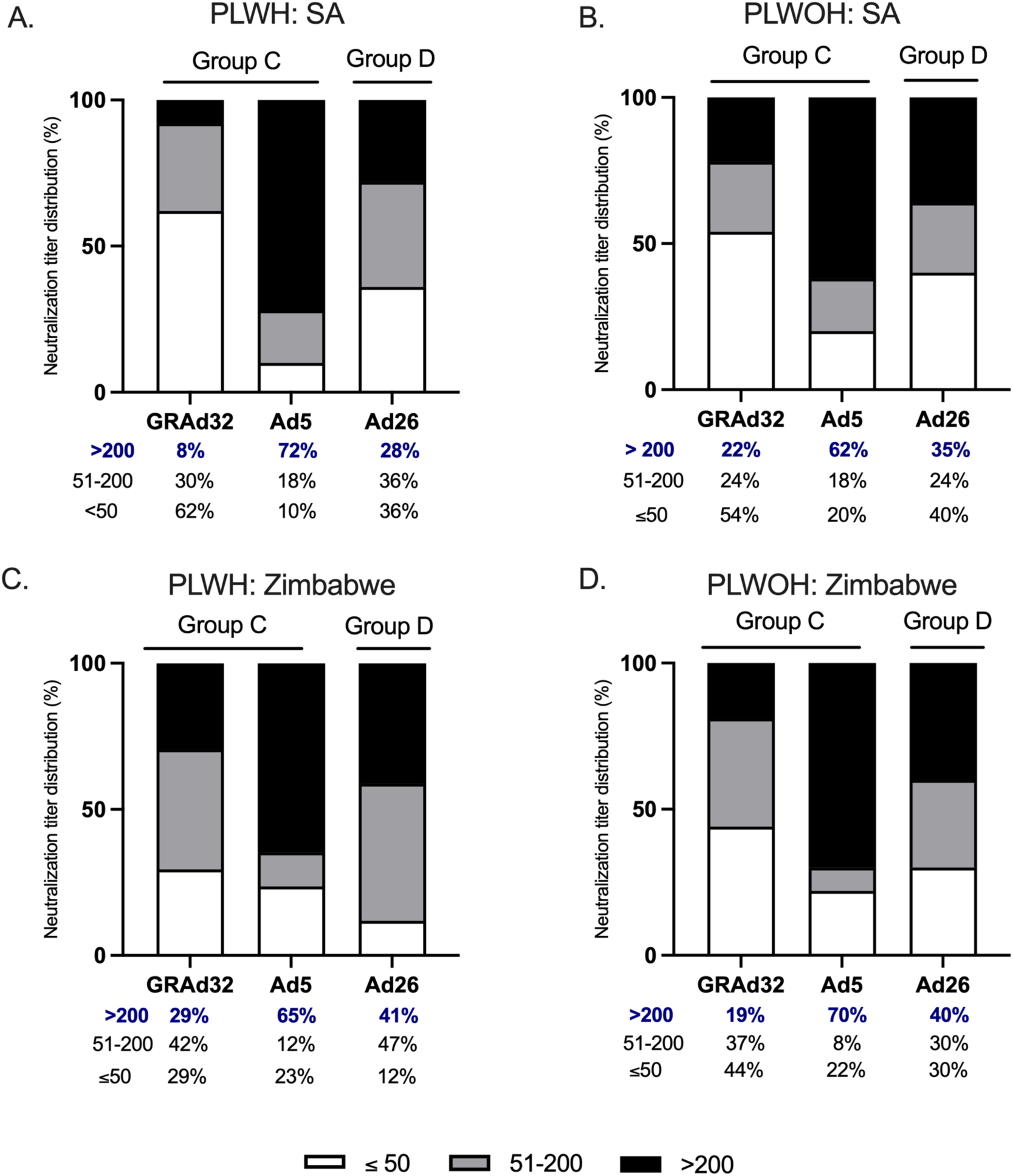
Titers to GRAd32 and other clinically relevant adenoviruses are similar in unvaccinated PLWH and PLWOH. Ad neutralizing antibody responses in unvaccinated individuals were measured and the proportional distribution of titers compared between A) PLWH and B) PLWOH in South Africa. Panels C and D show distribution of Ad responses in PLWH and PLWOH, respectively in Zimbabwe. Proportions of titers <50 (white), titers between 50 and 200 (gray) and titers >200 (black) are shown, with percentages below the graphs.

## DISCUSSION

Adenovirus-vectored vaccines are an attractive platform because they induce strong immune responses and have been used successfully in clinical settings. In addition to their immunogenicity and efficacy, they have less stringent cold-chain requirements than some newer platforms, making them particularly useful in resource-limited settings. A key consideration, however, is that pre-existing immunity to the vector can reduce immunogenicity to the vaccine, limiting its performance in populations with high background exposure to the vector. Although previous studies have assessed the prevalence of nAbs to GRAd32 in small cohorts from the United States of America and in Italy (Capone et al, 2021; Lanini et al, 2022), no prior studies had evaluated these responses in Africa. Therefore, in preparation for testing the GRAdHIVNE1 vaccine in South Africa and Zimbabwe (Paalangara et al. 2026)), we assessed the population-level neutralizing antibodies to GRAd32 (the GRAdHIVNE1 vector), as well as to other clinically relevant adenoviruses. Our results show that, while neutralizing antibodies to Ad5 and Ad26 are common, cross-reactive nAbs to GRAd32 are rare and generally of low titer. This remains true even after vaccination with an Ad26-based SARS-CoV-2 vaccine, in both PLWH and PLWOH. Overall, these findings suggest that the immunogenicity of GRAdHIVNE1 is unlikely to be significantly affected by pre-existing (natural infection or vaccine-induced) anti-adenovirus immunity, supporting the evaluation of this vaccine in sub-Saharan Africa.

Neutralizing antibodies to commonly circulating adenoviruses generally occur at higher levels in low medium income countries (LMICs), compared to high income countries (HICs). For example, high Ad5 seroprevalence (>70%) occurs in Africa, Asia, and South America, but is lower in Europe and North America (Barouch et al. 2011; Fuchs et al. 2015; Mennechet et al. 2019). Similarly, Ad26 antibodies were detected in <10% of adults in developed regions but in 31% and 66% of adults in Brazil and South Africa, respectively (Barouch et al. 2011; Fuchs et al. 2015)). These data raised concerns that even for less common non-human adenoviruses such as GRAd32, where low seroprevalence was reported in the USA and Italy, this may not be generalizable to African settings where seroprevalence maybe higher in these populations. However, our data suggests that seroprevalence of GRAd in South Africa and Zimbabwe is not very different to the US and Italy, which is in line with previous observations for other simian Ads (Quinn et al. 2013). This is reassuring for the ongoing clinical evaluation of GRAdHIVNE1. Future GRAd seroprevalence studies will need to expand to the rest of the African continent.

Although the GRAdHIVNE1 vaccine is being tested initially for HIV prevention, there is potential utility for this vaccine in PLWH, who might benefit from boosted CD8 T cell responses to conserved epitopes (Gaiha et al. 2019). Previous studies have indicated that PLWH, particularly those who are viremic and have impaired CD4 counts, may mount sub-optimal antibody responses to vaccines and infections (Motsoeneng et al. 2024). Indeed, PLWH show impaired immune responses to various respiratory infections, and often mount suboptimal responses to vaccines against viruses such as influenza, pneumococcus, hepatitis, and human papillomavirus (Tebas et al. 2010; Lacey 2019; Lee et al. 2014; Morsica et al. 2017). As southern Africa is still the epicenter of the HIV pandemic, we compared anti-Ad titers in PLWH and PWOH (Gengiah and Abdool Karim 2025). Overall, we saw no significant differences in viremically suppressed PLWH, accessing antiretroviral therapy. In ongoing studies, we will assess this in virally unsuppressed PLWH, who account for a quarter of PLWH in South Africa. However, these data suggest that GRAdHIVNE1 and other GRAd32-based vaccines could be deployed in PLWH.

We also assessed the impact of COVD-19 vaccination on titers of neutralizing antibodies. Administration of an Ad-based COVID-19 vaccine, Ad26.COV2.S, in the Sisonke cohort resulted in a significant boost in anti-Ad26 titers, even 9 months post-vaccination. This is consistent with other reports of increased vector immunity following homologous Ad boosting (Stephenson et al. 2021; Byazrova et al. 2022). We also observed a small increase in Ad5-directed titers following vaccination with Ad26.COV2.S (but not with inactivated vaccines). Ad26 belongs to species D and is phylogenetically distant from Ad5, with substantial divergence in hexon HVRs and fiber knob sequences, and limited antibody cross-reactivity reported (Mendonça et al. 2021). Moreover, if Ad26 vaccination indeed triggered increased Ad5 titers, one might expect these cross-reactive responses also to target GRAd32, which like Ad5, belongs to species C, but no increase in GRAd32 titers was observed. This aligns with results observed after vaccination with Ad4/Ad7 which showed limited cross-reactivity to Ad5 and select other Ads, rather than broad species cross-reactivity (Paris et al, 2014). Future monoclonal antibody isolation studies in these participants may provide insights into cross-reactivity.

The impact of anti-vector neutralization remains unclear. Several studies showed that repeat administration of Ad26.COV2.S resulted in boosting of anti-spike titers (Sadoff et al. 2022; Le Gars et al. 2022; He et al. 2022; Moyo-Gwete et al. 2023). In the BaSiS cohort, a homologous Ad26.COV2.S vaccine failed to boost anti-spike titers, unlike the Pfizer arm, though this was not directly attributed to anti-vector immunity (Riou et al. 2024). Similarly, studies of adenovirus-vectored HIV and Ebola vaccines have mixed conclusions. Analysis of the HIV STEP trial found that high pre-existing Ad5 neutralizing antibody titers were associated with reduced immunogenicity (Cheng et al. 2012) and the same was reported for the Ad5-based COVID-19 vaccine from CanSino (**Zhu et al 2020)**. In contrast, studies of Ad26.ZEBOV (Barry et al. 2021), ChAdOx1 nCoV-19 (Madhi et al. 2022; Ogbe et al. 2021) and ChAd3 (Ewer et al. 2016) found limited impact of pre-existing neutralizing antibodies on immunogenicity of the transgene. The clinical trial of GRAdHIVNE1 will provide an opportunity to comprehensively assess the impact of anti-GRAd32 neutralizing antibodies on HIV-1 vaccine immunogenicity.

Limitations of this study include that only neutralizing antibody responses were measured, and anti-vector T cell responses were not assessed, despite their potential to modulate the immunogenicity of adenovirus-vectored vaccines. In addition, comparisons between Ad26-vaccinated and non-Ad vaccine (inactivated) recipients were made across distinct cohorts from different countries, which may introduce unmeasured confounding. Finally, limited information was available for the Zimbabwean cohort, regarding the timing of vaccination, constraining interpretation of those findings. Despite these limitations, these findings indicate that anti-vector antibody responses are unlikely to compromise the immunogenicity of GRAdHIVNE1, supporting its evaluation and even future deployment in sub-Saharan Africa.

## Acknowledgements.

The authors thank the participants, site Principal investigators, laboratory staff and protocol-PIs of the Sisonke, BaSiS and Mutala cohort studies. We thank S. Barnett and T. Onami, senior program officers from the Gates Foundation, for their guidance and support.

## Disclaimer

The analyses, conclusions, opinions, and statements expressed herein are solely those of the authors and do not reflect those of the funding or data sources; no endorsement is intended or should be inferred.

## Data Availability

All data is available upon reasonable request.

