## Supplemental Figure 1 for "Low seroprevalence of neutralizing antibodies to gorilla adenovirus 32 (GRAd32) in southern African populations supports evaluation of this vector platform for HIV vaccine development"

***S1 Fig: Neutralizing antibodies to GRAd32 are rare in unvaccinated South African and Zimbabwean adults, and comparable to US adults***

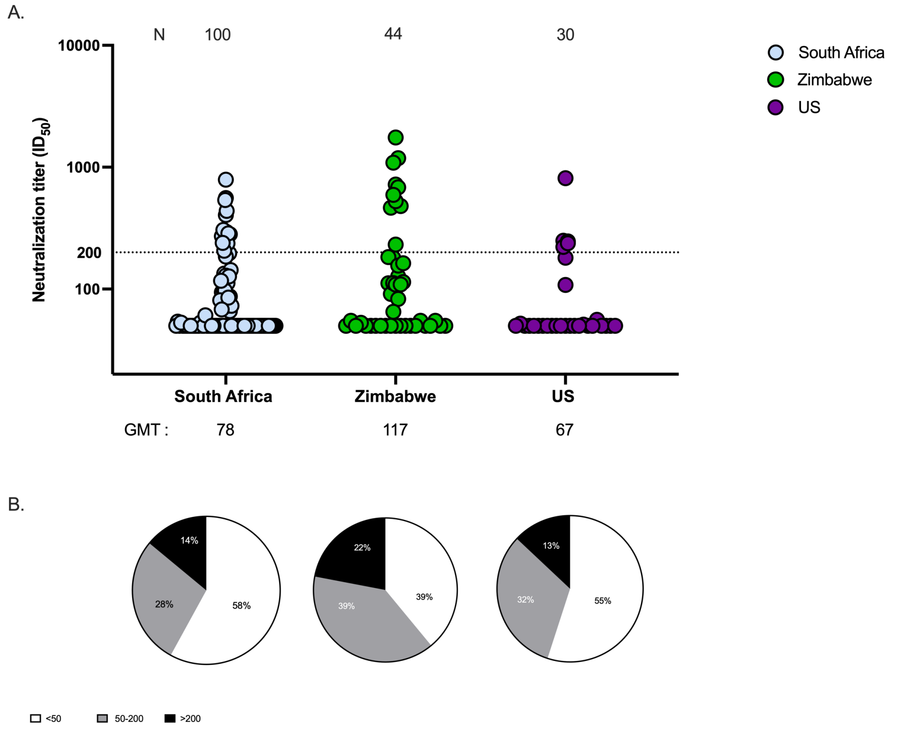


***S1 Fig: Neutralizing antibodies to GRAd32 are rare in unvaccinated South African and Zimbabwean adults, and comparable to US adults.*** *Anti-GRAd32 neutralization responses across South Africa (blue dots), Zimbabwe (green dots) and the US (purple dots) were measured. Geometric mean titers (GMT) are shown below the graph. (B) Anti-GRAd32 neutralization antibody titer distribution pie-charts across the different cohorts, with proportions shown as titers <50 (white), titers 50-200 (gray) and titers >200 (black).*
