## Supplemental Figure 2 for "Low seroprevalence of neutralizing antibodies to gorilla adenovirus 32 (GRAd32) in southern African populations supports evaluation of this vector platform for HIV vaccine development"

***S2 Fig: Ad26.COV2.S vaccination results in increased titers to Ad26, but does not impact GRAd32 titers.***

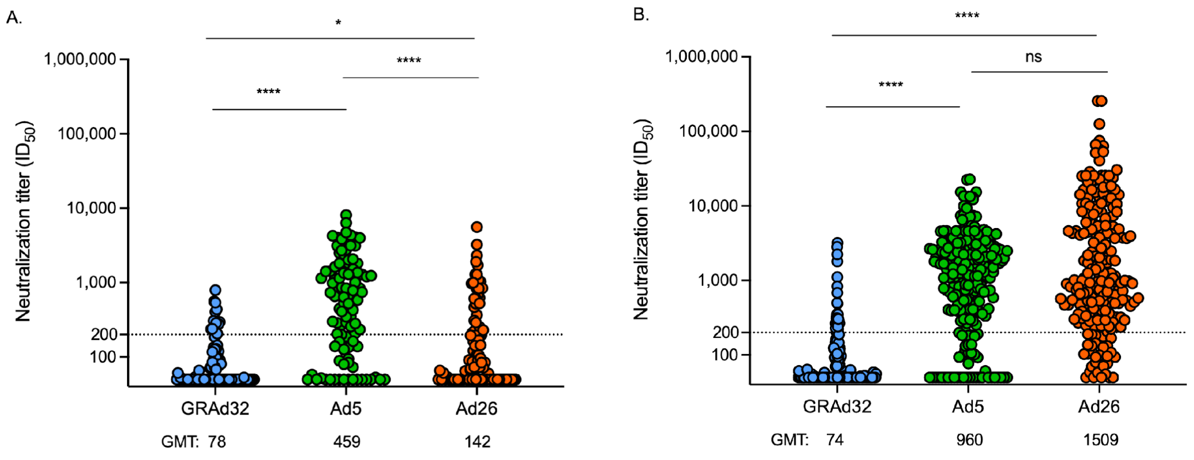


***S2 Fig: Ad26.COV2.S vaccination results in increased titers to Ad26, but does not impact GRAd32 titers.*** *Serum samples from participants enrolled in the Sisonke and BaSiS baseline timepoints, ie before Ad26.COV2.S vaccination (n=100) (A) and 9 months after Ad26.COV2.S vaccination (n=226) (B) were tested for Ad nAb responses to GRAd32 (blue dots), Ad5 (green dots) and Ad26 (orange dots). Each dot represents a single participant. Geometric mean titers (GMT) are shown by horizontal lines, and values are shown below the graph. Dotted lines indicate titers of 200 (titer associated with reduced immunogenicity for vaccines).*
