## Supplemental Figure 3 for "Low seroprevalence of neutralizing antibodies to gorilla adenovirus 32 (GRAd32) in southern African populations supports evaluation of this vector platform for HIV vaccine development"

**S3 Fig: Neutralizing antibodies to adenoviruses are similar in unvaccinated Zimbabwean adults and those vaccinated with inactivated COVID vaccines**

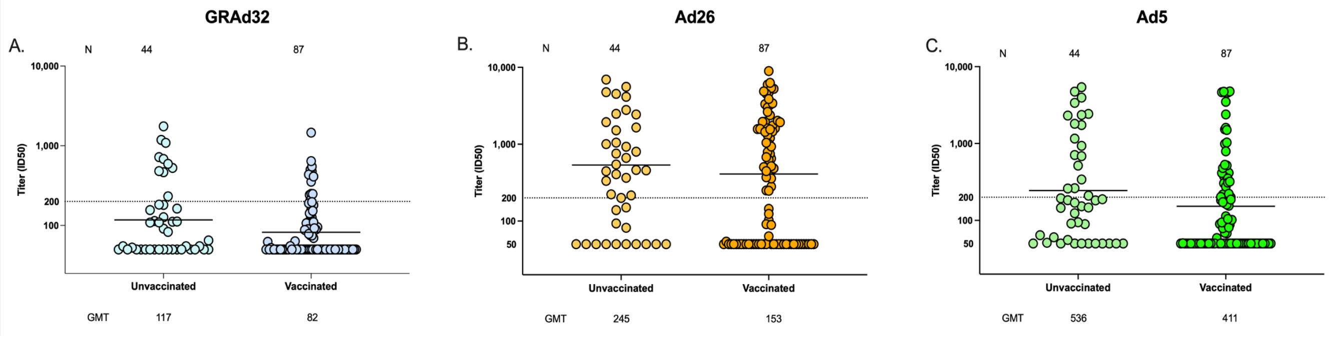


**S3 Fig: Neutralizing antibodies to adenoviruses are similar in unvaccinated Zimbabwean adults and those vaccinated with inactivated COVID vaccines.** *Neutralization responses to GRAd32 (A), Ad26 (B) and Ad5 (C) in SARS-CoV-2 vaccinated and unvaccinated adults from Zimbabwe, with each dot representing a single participant. Geometric mean titers (GMT) are shown by horizontal lines, and values are shown below the graph. Dotted lines indicate titers of 200 (titer associated with reduced immunogenicity for vaccines).*
